# pH-dependent polymorphism of the structure of SARS-CoV-2 nsp7

**DOI:** 10.1101/2021.09.10.459800

**Authors:** Yeongjoon Lee, Marco Tonelli, Mehdi Rahimi, Thomas K. Anderson, Robert N. Kirchdoerfer, Katherine Henzler-Wildman, Woonghee Lee

**Affiliations:** Department of Chemistry, University of Colorado Denver, Denver, CO 80204, USA; National Magnetic Resonance Facility at Madison (NMRFAM), University of Wisconsin at Madison, Madison, WI 53706, USA; Biochemistry Department, University of Wisconsin-Madison, Madison, WI 53706, USA; Institute for Molecular Virology, University of Wisconsin-Madison, Madison, WI 53706, USA

## Abstract

The solution structure of SARS-CoV-2 nonstructural protein 7 (nsp7) at pH 7.0 has been determined by NMR spectroscopy. nsp7 is conserved in the *coronavirinae* subfamily and is an essential co-factor of the viral RNA-dependent RNA polymerase for active and processive replication. Similar to the previously deposited structures of SARS-CoV-1 nsp7 at acidic and basic conditions, SARS-CoV-2 nsp7 has a helical bundle folding at neutral pH. Remarkably, the α4 helix shows gradual dislocation from the core α2-α3 structure as pH increases from 6.5 to 7.5. The protonation state of residue H36 contributes to the change of nsp7’s intramolecular interactions, and thus, to the structural variation near-neutral pH. Spin-relaxation results revealed that all three loop regions in nsp7 possess dynamic properties associated with this structural variation.

## Introduction

Since 2002, coronaviruses (CoVs) have threatened human health with three outbreaks: severe acute respiratory syndrome (SARS), Middle Eastern respiratory syndrome (MERS), and the coronavirus disease of 2019 (COVID-19).^1^ In just a year after COVID-19 was declared a global pandemic by the WHO in March 2020, more than 119 million individuals have been infected with SARS-CoV-2, with 2.6 million deaths across the world. To address the COVID-19 pandemic, research efforts have led to new advances in the basic understanding of SARS-CoV-2. Similar to the other highly pathogenic CoVs, SARS-CoV-2 belongs to the genus *Betacoronavirus*.^2^ The genome of SARS-CoV-2 shares high sequence identity with those of SARS-CoV-1 (∼80 %) and MERS-CoV (∼50 %).^3,4^ The genome consists of 14 open reading frames (ORFs), with the two largest encoding two large polyproteins, that once cleaved, comprise the 16 viral non-structural proteins or nsps.^4^ The remaining open reading frames encode the four structural proteins – spike (S), envelope (E), membrane (M), and nucleocapsid (N) – and nine accessory proteins.^4^

SARS-CoV-2 is an enveloped, positive-sense RNA virus.^2^ Upon entering host cells, the two largest and 5’ proximal ORFs are translated into polyproteins that are subsequently cleaved into individual nsps.^1,5^ The mature nsps induce rearrangement of the endoplasmic reticulum (ER) into double-membrane vesicles (DMVs) that are platforms for viral replication of genomic and subgenomic RNAs.^1,6,7^ Translation of the subgenomic mRNA’s produces structural and accessory proteins.^8^ These proteins and the newly synthesized genomic RNA assemble into a virus particle in the ER-Golgi intermediate compartment (ERGIC) that is later secreted to the intercellular space by the secretory pathway.^1,9^

nsp7, 8, and 12 are incorporated into DMVs and form the core structure of the viral replication-transcription complex (RTC).^5,7,10^ These core proteins are highly conserved in coronaviruses.^11-13^ While nsp12 serves as the catalytic center being the RNA-dependent RNA Polymerase, nsp7 and nsp8 are essential co-factors for processive RNA synthesis.^14-16^ To examine the crucial roles in viral replication and transcription of these protein in the virual life cyle, many crystallographic and electron microscopic structures have been determined. By contrast, the structural and dynamic features of nsps in solution have not been as widely investigated. Here, we determined the solution structure of SARS-CoV-2 nsp7 at pH 7.0 and compared it with structures of SARS-CoV-1 nsp7 at different pHs to better understand its structure in the biophysiological environment. We found that nsp7 shows gradual changes in its morphology as the pH changes. The pH-dependent protonation state of H36 (native number) was found to be a key element that causes this structural variation. The motional properties of the backbone of nsp7 were investigated. Our findings may provide an essential clue for understanding the structural and dynamic features of nsp7 associated with cellular pH conditions.

## Results and Discussion

### Solution Structure of SARS-CoV-2 nsp7 at pH 7.0

Similar to the structure of SARS-CoV-1 nsp7 at pH 7.5 and 6.5,^17,18^ SARS-CoV-2 nsp7 exhibited a helical bundle structure at pH 7.0 (Fig. 1a). Four helices defined by PROCHECK^19^ – α1 (V11-L20), α2 (K27-L41), α3 (T45-M62), and α4 (I68-R79) – are aligned with the TALOS-N prediction (green regions) reported in a previous paper.^20^ The statistics for the final 20 lowest energy models are shown in Table 1. Only the backbone and heavy atoms of residues in the ordered region (S10–R79) were included in the calculation of RMSDs. Except for the highly disordered N-terminal regions, the ordered regions of all calculated models were well-superimposed (Fig. 1b).

**Table 1.**
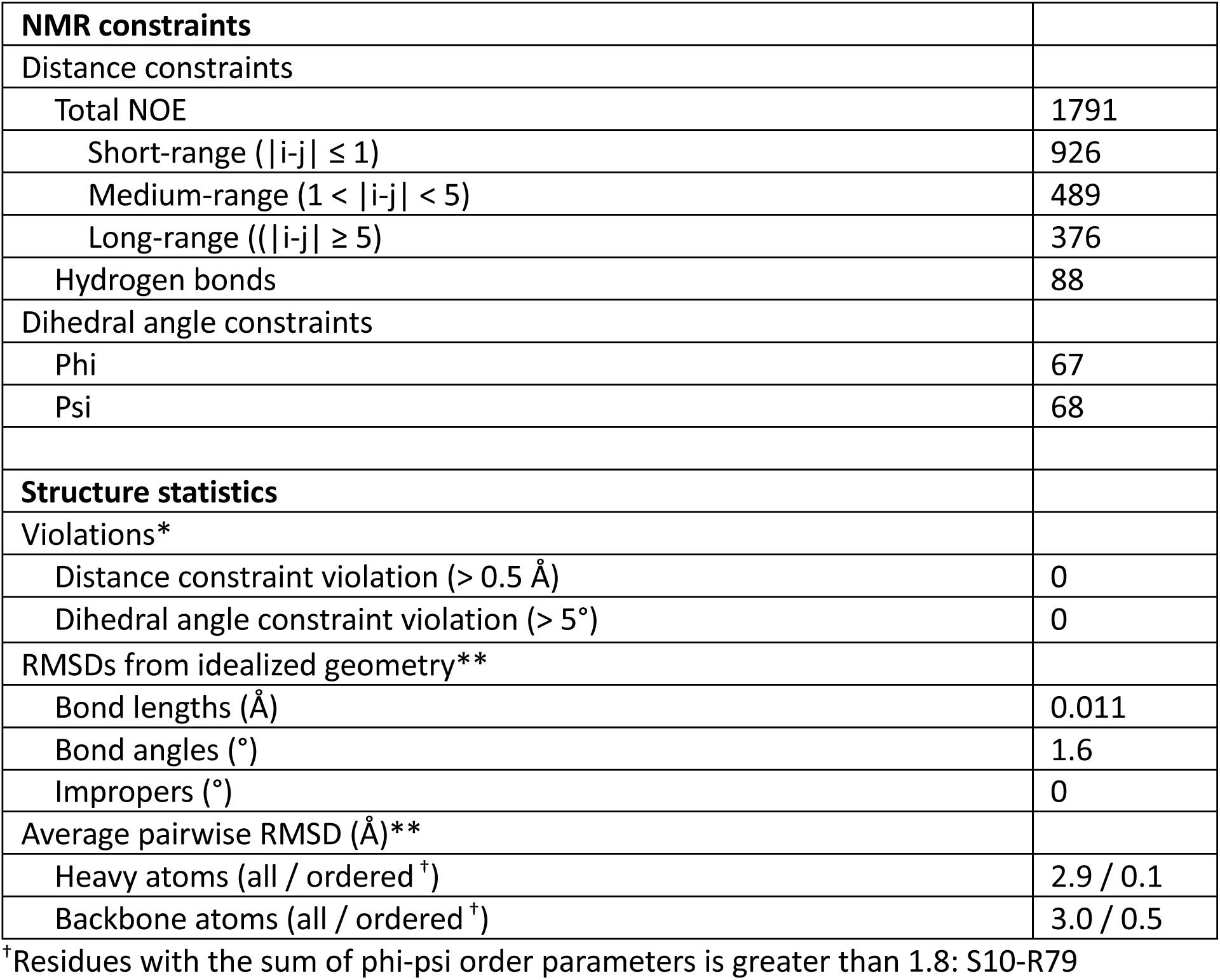
NMR and refinement statistics for the solution structure of SARS-CoV-2 nsp7 at pH 7.0. The 20 lowest energy models were evaluated with *Poky Analyzer^37^ and **PSVS 1.5.^39^

**Figure 1.**
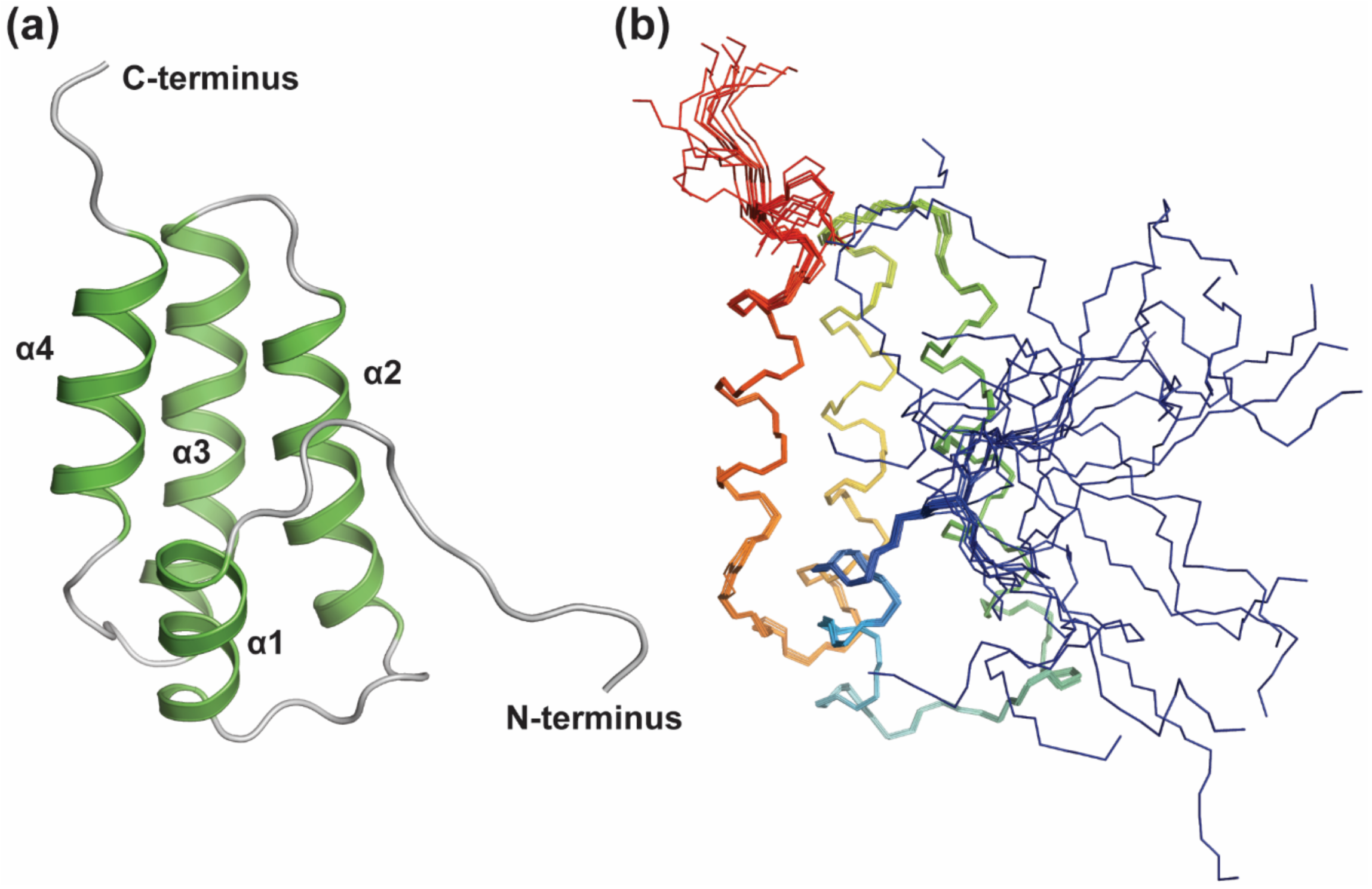
Structural features of SARS-CoV-2 nsp7 at pH 7.0. (a) Four helices of SARS-CoV-2 nsp7 align well with the TALOS-N prediction (green)^20^ and form a helical bundle structure that comprises four helices. (b) Backbone ensemble structure of the 20 lowest energy models.

### pH-dependent Polymorphism of the Structure of Coronavirus nsp7

Since there are no determined structures for SARS-CoV-2 nsp7 at different pH values, the structures of SARS-CoV-1 nsp7 at pH 7.5 (PDB ID 1YSY)^17^ and 6.5 (PDB ID 2KYS)^18^ were used to investigate the pH-dependency of the SARS-CoV-2 nsp7 structure. Given the high degree of sequence and structural homology, we compared the structures of SARS-CoV-1 at pH 6.5 and 7.5 together with our structure of SARS-CoV-2 at pH 7 to assess the pH-dependent polymorphism of nsp7. As shown in Fig. 2a, SARS-CoV-1 and SARS-CoV-2 nsp7 differ by a single residue at position 70 in the α4 helix, where SARS-CoV-2 has a Lys rather than an Arg. However, since the sidechains of both Lys and Arg in both nsp7s do not contact any residue on the other helices, they are irrelevant to the tertiary folding (Fig. 2b-d). Thus, the conformations of SARS-CoV-2 nsp7 at pH 7.5 and 6.5 are expected to be similar to those of SARS-CoV-1 nsp7 at the corresponding pHs.

**Figure 2.**
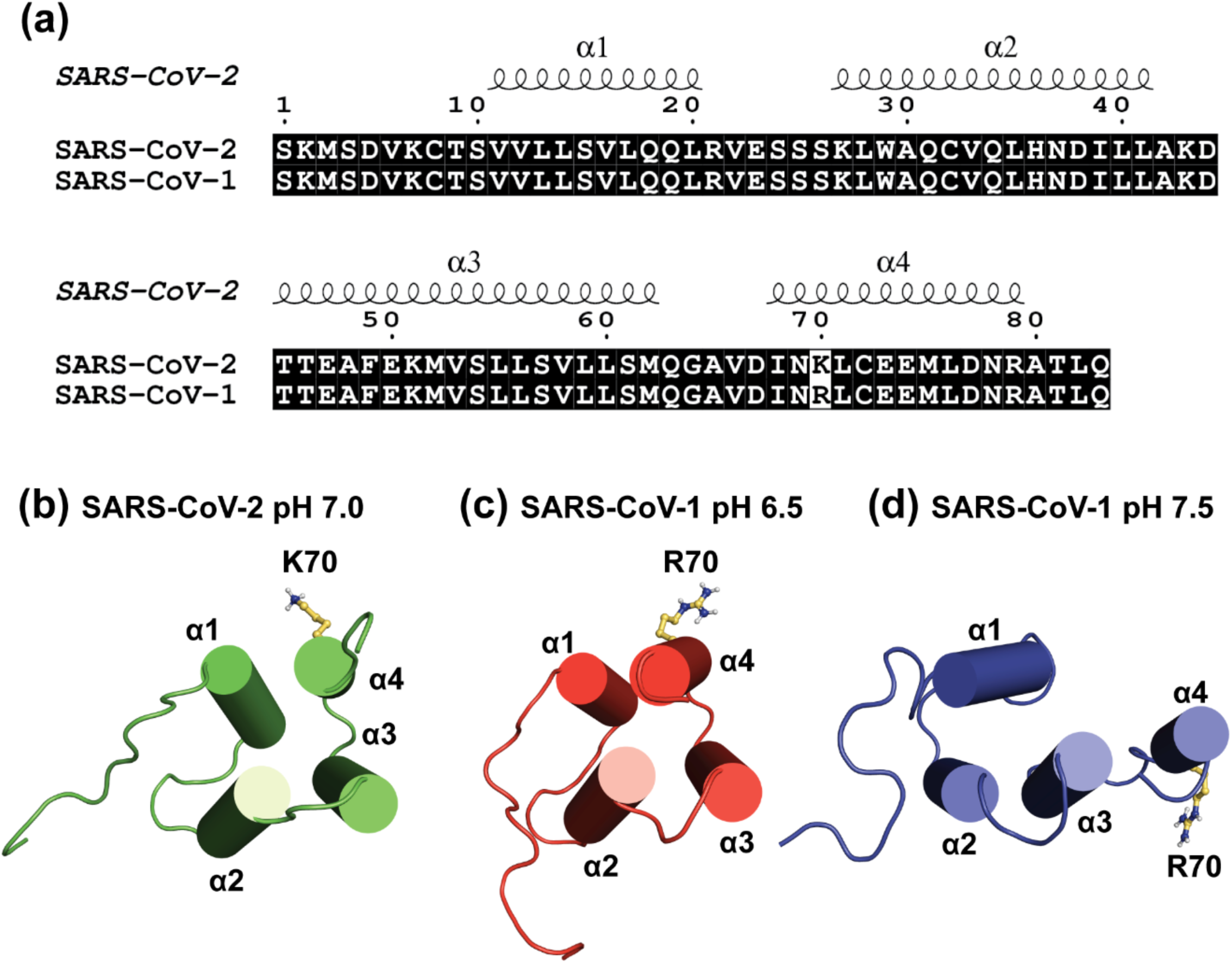
Comparison of the primary and tertiary structures of SARS-CoV nsp7. (a) Sequence alignment of nsp7 from two sarbecoviruses. Four spirals indicate the helical regions of SARS-CoV-2 nsp7 at pH 7.0. Identical residues are highlighted black. The only difference between the two protein sequences is the type of basic residue at the N-terminal part of α4 helix. (b) The sidechain of K70 in SARS-CoV-2 nsp7 at pH 7.0 has no interaction with residues in other regions. The same feature is also shown for R70 in SARS-CoV-1 at (c) pH 6.5 and (d) pH 7.5. Although the overall folding of nsp7 differs greatly with pH, it does not appear to be affected by the difference in one basic residue.

The structures were superimposed based on the core structure of the protein that comprises the α2 and α3 helices (Fig. 3). As observed previously, the core structure of nsp7 is almost independent of pH changes within the 6.5 to 7.5 range.^18^ However, the α4 helix is gradually dislocated from the core structure as pH increases, completely losing contact with the α2 helix at pH 7.5. Since the α4 helix is known as the stabilizing factor for the overall folding,^18^ this dislocation may destabilize the structure of nsp7. This is compensated by the translocation of the α1 helix. As the α4 helix is displaced, the α1 helix moves to the position originally occupied by the α1 and α4 helices (Fig. 3b). By forming new interactions with the core structure, the α1 helix can act as a supplementary factor for maintaining the overall folding at pH 7.5.

**Figure 3.**
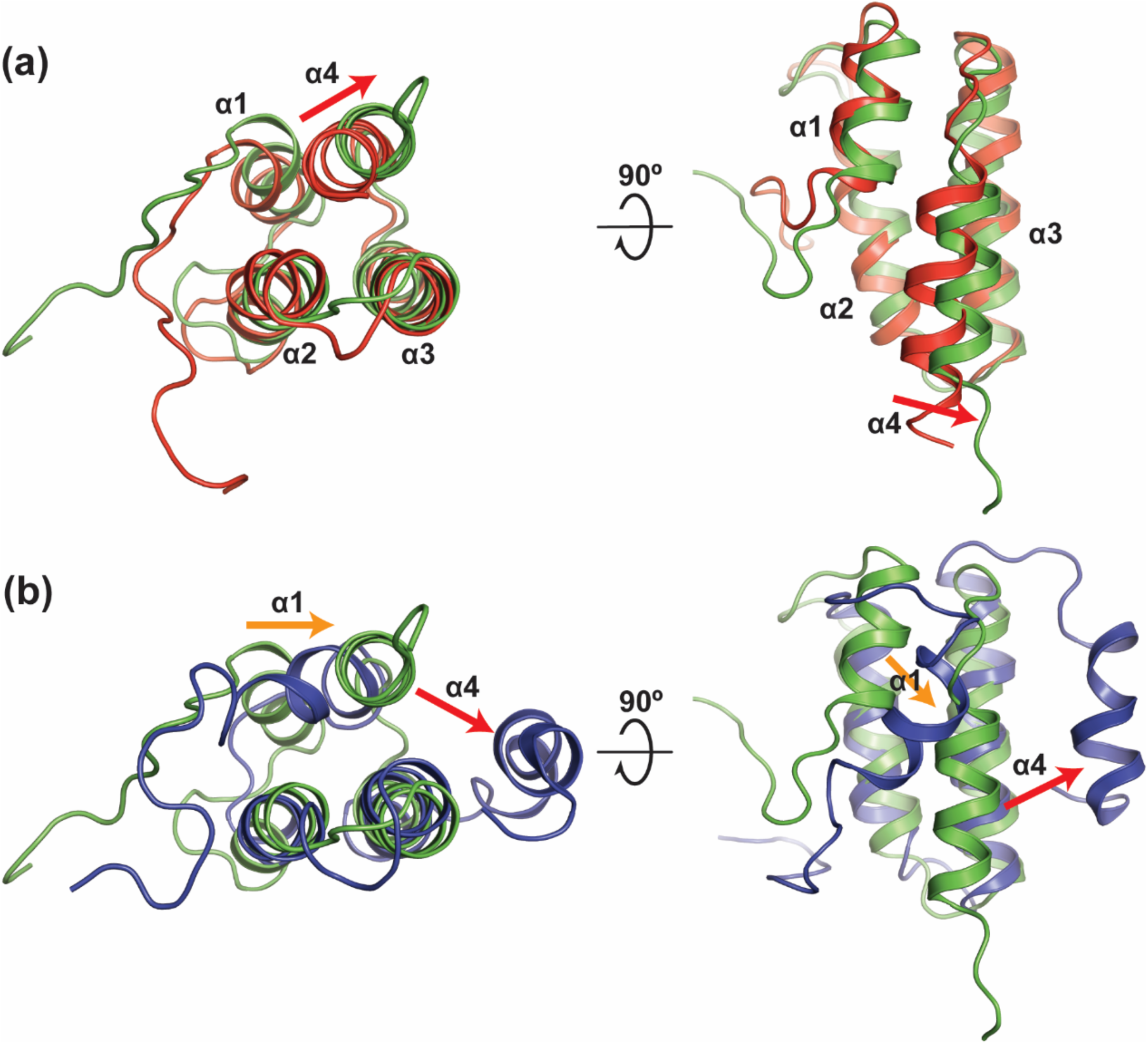
pH-dependent polymorphism of nsp7 structure. (a) Structural alignment of SARS-CoV-1 nsp7 at pH 6.5 (red) with SARS-CoV-2 nsp7 at pH 7.0 (green). As the pH increases the α4 helix becomes slightly dislocated from the core α2-α3 structure. (b) Structural alignment of SARS-CoV-2 nsp7 at pH 7.0 (green) and SARS-CoV-1 at pH 7.5 (blue). At the weakly basic pH, the α4 helix becomes totally displaced and the α1 helix moves to the place originally occupied by the α1 and α4 helices at the neutral pH.

### Protonation State of a Histidine Residue and pH-dependent Polymorphism

Since the sidechain of histidine residues have a pK_a_ value near neutral pH, its protonation states are often closely related to the pH-dependent variations of protein structure and oligomerization.^21-26^ Accordingly, the observed pH-dependent polymorphism of the nsp7 structure seems to be related to the protonation state of H36 in the α2 helix. H36 is conserved in both SARS-CoV-1 and SARS-CoV-2 nsp7. At acidic pH, the Nε2 atom of H36 prefers to be protonated. The Hε2 atom can then form a hydrogen bond with the Oδ1 atom of N37 (Fig. 4a). On the opposite side of the His ring, the Nδ1-Hδ1 bond serves as a proton donor for another hydrogen bond with the E74 from the α4 helix. Along with the hydrophobic packing between α2-α3-α4 helices, these hydrogen bonds may contribute to the compact fold of nsp7 at acidic pH, with the α4 helix tightly binding to the core structure of the protein.

**Figure 4.**
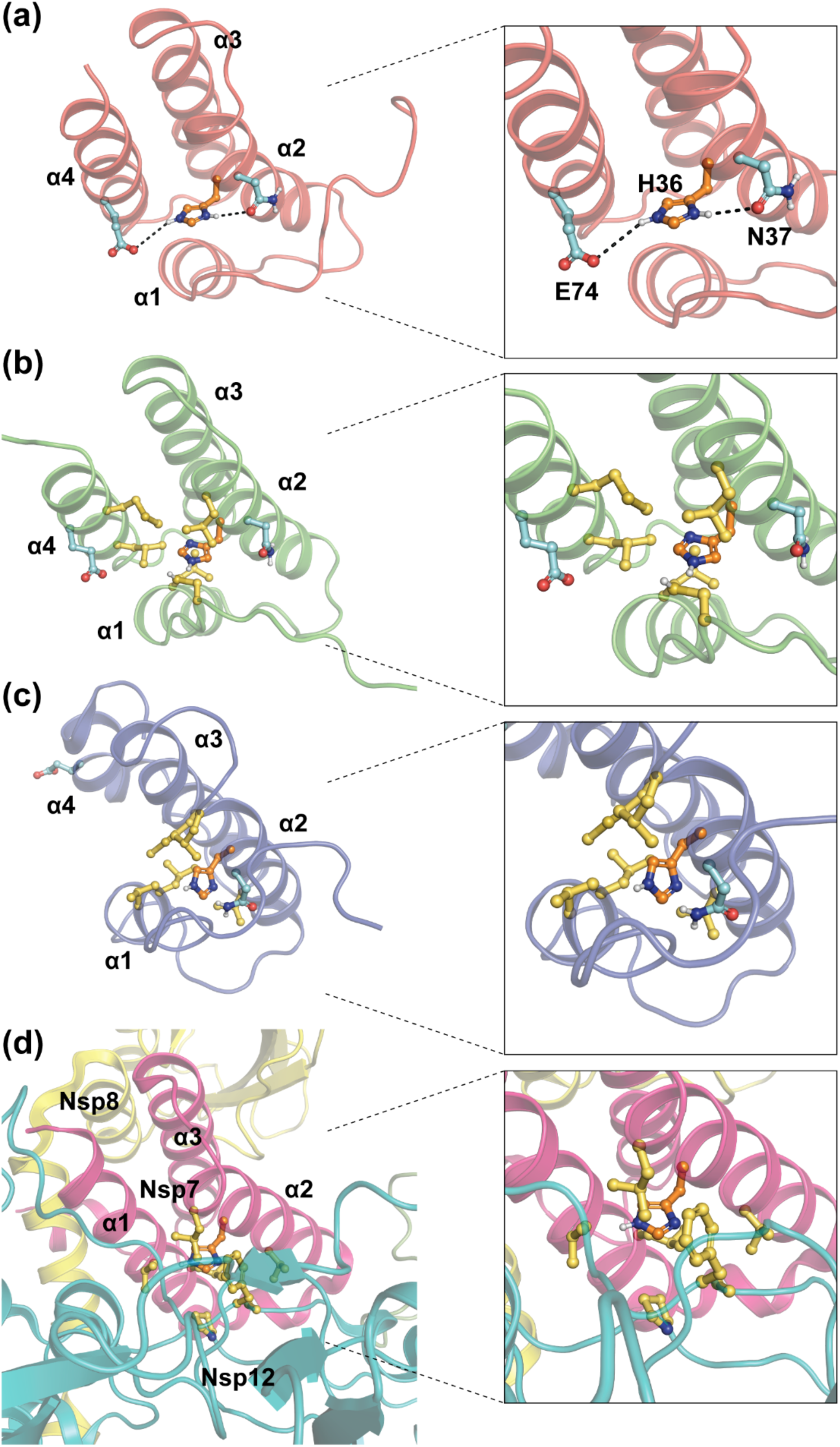
H36 and the structural variations of nsp7. The sidechains of H36 and its interacting residues are shown in various conditions. (a) In the aqueous solution at pH 6.5, the protonated sidechain of H36 (orange) forms two hydrogen bonds with the sidechains of N37 and E74 (cyan), thus contributing to the compact conformation of nsp7. (b) As the sidechain of H36 becomes deprotonated at pH 7.0, it loses the hydrogen bonds and becomes surrounded by hydrophobic residues (yellow) from α1-α2-α4 helices. (c) As the α4 helix is displaced at pH 7.5, the sidechain of H36 forms a new hydrophobic packing between α1-α2-α3 helices. (d) The sidechain of H36 also exists in the deprotonated state in the crystal structure of the replication complex, where it participates in the hydrophobic interactions between nsp7 and Nsp12.

As the pH increases to 7.0, the population of deprotonated H36 significantly increases, leading to the loss of the electrostatic interaction. This in turn leads to a conformational rearrangement that dislocates the α4 helix slightly outward from the core structure. The neutralized aromatic sidechain is now slightly buried within the hydrophobic packing among α1-α2-α4 helices (Fig. 4b), leading to a large decrease of solvent-accessible surface area (SASA) from 16.133 Å^2^ to 1.230 Å^2^. Nonetheless, as H36 balances between protonated and deprotonated forms, the overall structure of nsp7 still resembles the compact conformation. However, as the pH further increases above 7.0, the majority of H36 becomes deprotonated and the structure of nsp7 experiences a dramatic conformational change (Fig. 4c). The completely dislocated α4 helix no longer makes contact with the α2 helix, and only barely contacts the α3 helix by a few hydrophobic interactions. As the α1 helix translocates to occupy the position vacated by the α4 helix, the sidechain of H36 is surrounded by N37 and hydrophobic residues from the α1, α2, and α3 helices, thus losing SASA (0 Å^2^) completely. Consequently, H36 participates in the newly formed hydrophobic packing that may contribute to maintain the folding of the core structure at basic pH.

The structural variations of proteins in different environments often lead to different intermolecular interactions and may have a profound effect on the assembly state of protein complexes. Thus, given the observed effect of pH on the structure of nsp7, we propose that the pH-dependent polymorphism of nsp7 may play a critical role in the assembly state of the overall replication complex. During viral infection, nsp7 experiences significant pH changes as it travels between different cellular compartments. Immediately after production, nsp7 would exist as a globular protein that reversibly exchanges between two conformations, a more compact one, preferred in the acidic condition, and a loose one, preferred in the basic condition.^18^ At the slightly basic pH of DMV (pH 7.0 – 7.2),^27^ nsp7 in its loose conformation associates with its binding partners, Nsp8 and 12, to form the replication complex. The N-terminal region of nsp7 becomes fully structured, and the extended α1 helix forms a three-helix bundle with the core α2-α3 structure.^11-13^ In this condition, the sidechain of H36 is packed within the hydrophobic residues at the binding interfaces of nsp7 and nsp12, as illustrated in Fig. 4d. If the sidechain of H36 becomes protonated, its positive charge would make the hydrophobic interactions between nsp7 and nsp12 unfavorable. In this way, the protonation state of H36 of nsp7 would affect the assembly status of RTC, and thus, the viral replication.

### Spin-relaxation Experiment of SARS-CoV-2 nsp7 at pH 7.0

Previously, the line shape analysis for the ^1^H-^15^N HSQC peaks of SARS-CoV-1 nsp7 revealed that the protein may have a dynamic loop between α3 and α4 helices at pH 6.5. Residues near the C-terminal end of α3 helix (S61 and Q63), in the loop 3 (V66), and near the N-terminal end of α4 helix (N69 and L71) exhibited a noticeable line-broadening that may be related to the conformational exchange of α4 helix at the weakly acidic pH.^18^ To understand the dynamic properties of SARS-CoV-2 nsp7 in detail, the longitudinal (R_1_) and transverse (R_2_) relaxations and heteronuclear Overhauser effect (hetNOE) experiments were conducted at the neutral pH (Fig. 5 and 6). Our results confirmed that all three loop regions of SARS-CoV-2 nsp7 have dynamic features at pH 7.0 that are needed for the exchange between compact and loose conformations.

**Figure 5.**
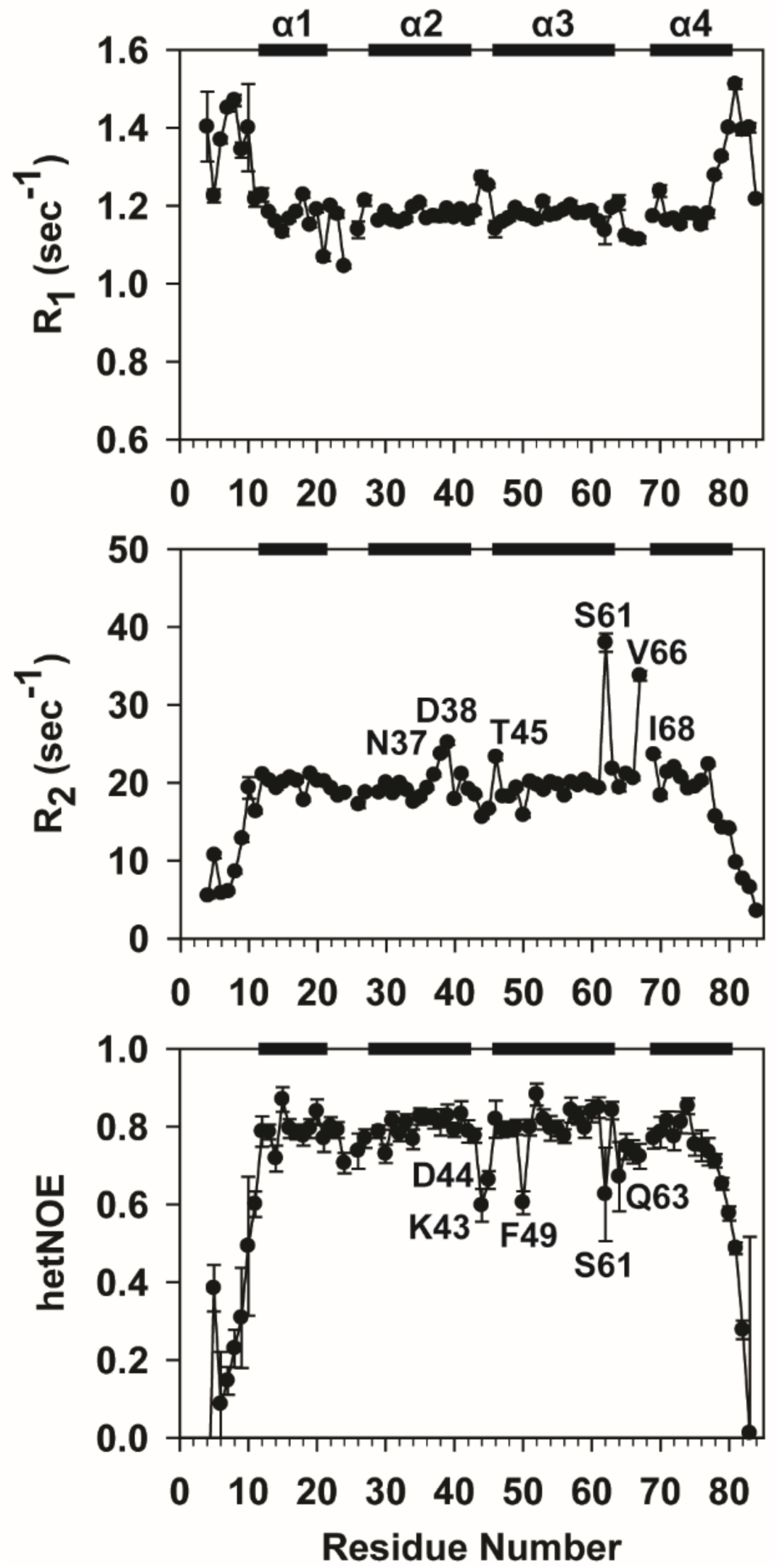
Spin-relaxation results of SARS-CoV-2 nsp7 at pH 7.0. It was not able to determine R_1_, R_2_, and hetNOE values for S24, K27, and D67 because their peaks were not seen in the spectra. The secondary structures are shown as black bars at the top of each plot.

**Figure 6.**
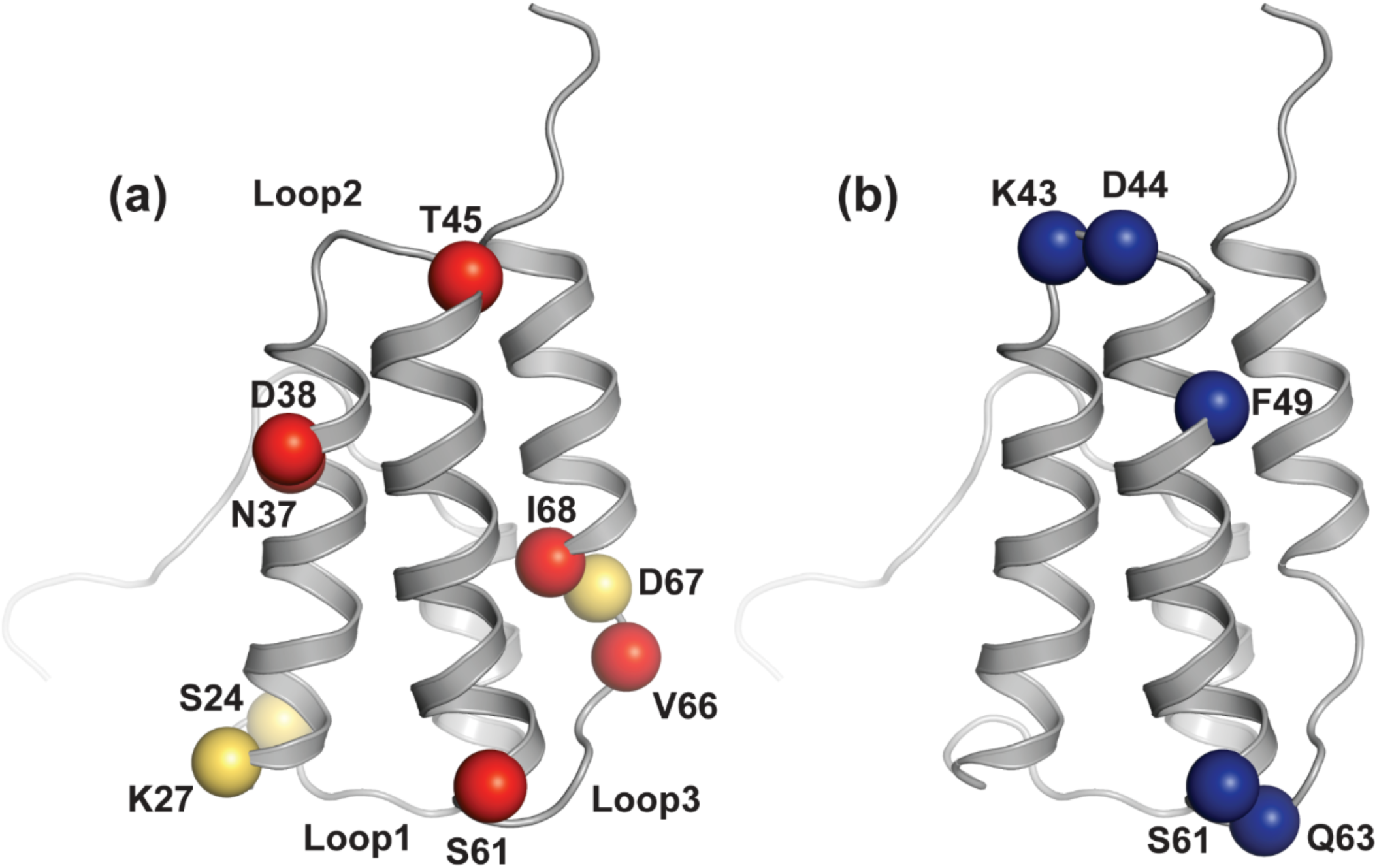
Visualization of spin-relaxation results on the nsp7 structure. (a) The alpha carbon of residues with R_2_ higher than 23 sec^-1^ are shown as red spheres. Also, S24, K27, and D67 whose peaks are not found in the spectra are shown as yellow spheres. (b) Residues with hetNOE lower than 0.7 are shown as blue spheres. For better interpretation, residues in the highly flexible terminal regions are excluded from the visualization.

As expected, S61 near the C-terminal end of α3 helix, V66 in the loop 3, and I68 at the N-terminal end of α4 helix were found to have high R_2_ rates, 37.96 ± 1.20 sec^-1^, 33.72 ± 0.63 sec^-1^, and 23.60 ± 0.27 sec^-1^, respectively. It seems that a slight difference in the nsp7 structure at pH 6.5 and 7.0 changed the motional properties at Q63, I68, N69, and L71. Despite this difference, it is still clear that the broad region in between the ends of α3 and α4 helices has sub-nanosecond timescale motions that are related to the movement of α4 helix. Interestingly, the amide proton of D67 in this dynamic region seems to have a very short transverse relaxation time (T_2_) since its peak was not shown in any NMR spectrum probably due to line-broadening. Similarly, the peaks for S24 and K27 in the loop 1 showed very weak intensities in most NMR spectra and were not shown in the spin-relaxation experiments, indicating their short T_2_ times. Therefore, the backbone regions including these two residues also have dynamic motions related to the movement of α1 helix.

Similar to the previous hetNOE results for SARS-CoV-1 nsp7 at pH 6.5,^18^ our results for SARS-CoV-2 nsp7 at pH 7.0 also revealed the increased structural flexibility at the highly disordered N- and C-terminal regions. In addition, the protein revealed two highly flexible loop regions, loop 2 and loop 3. In loop 2, K43 and D44 showed low hetNOE values, 0.597 ± 0.042 and 0.662 ± 0.023, respectively. Although T45, located between the end of loop 2 and the beginning of helix α3, was found to have a high hetNOE, it also showed a high R_2_ rate, 23.28 ± 0.41 sec^-1^, suggesting the presence of sub-nanosecond chemical exchange at the end of loop 2. As for loop 3, Q63 was found to have the lowest hetNOE of 0.671 ± 0.089, with the remaining three residues G64, A65, and V66 also showing low hetNOE values close to 0.7.

Remarkably, N37 and D38 near H36 showed significantly increased R_2_ rates, 23.65 ± 0.22 sec^-1^ and 25.14 ± 0.26 sec^-1^, respectively. Although the reason is still unclear, it seems that the structural variations by the protonation/deprotonation of H36 are related to their sub-nanosecond motions. Also, we measured an unexpectedly low hetNOE value of 0.604 ± 0.030 for F49 in the α3 helix. It was found that the sidechain of F49 is exposed toward the solvent (SASA = 114.166 Å^2^). This hydrophobic sidechain may tremble in the solvent, making its backbone region unexpectedly flexible.

The correlation time (τ_c_) of the Brownian rotation diffusion of a small rigid protein in solution can be calculated as a function of the ratio of longitudinal and transverse relaxation times, T_1_ and T_2_. ^28,29^ Given that the τ_c_ values of globular proteins are proportional to their molecular weight (MW), one can use the calculated τ_c_ to approximate the MW of a protein using a standard curve of τ_c_ versus MW at a given temperature. ^28^ For SARS-CoV-2 nsp7 at pH 7.0 and 298 K, τ_c_ was calculated to 12.4 ns. Using the standard curve created with empirical data from NESG^28^ (Fig. 7), the MW of nsp7 was approximated to 20.6 kDa, which is about twice the value expected for the monomer (9.4 kDa). This suggests that SARS-CoV-2 nsp7 may exist as a homodimer when it is isolated in solution. Nevertheless, it is still unclear whether the protein really dimerized in the NMR sample, as we were unable to find any distinguishable intermolecular NOEs in the NOESY spectra. Accordingly, it was not possible to determine the dimeric structure because of the lack of contact information. We plan to verify whether nsp7 dimerizes in solution, and if it does, whether dimerization is a natural property of the protein or just an unintended consequence of the high concentration required for NMR studies. We also plan to verify whether the dimerization of nsp7 has any effect on its conformation.

**Figure 7.**
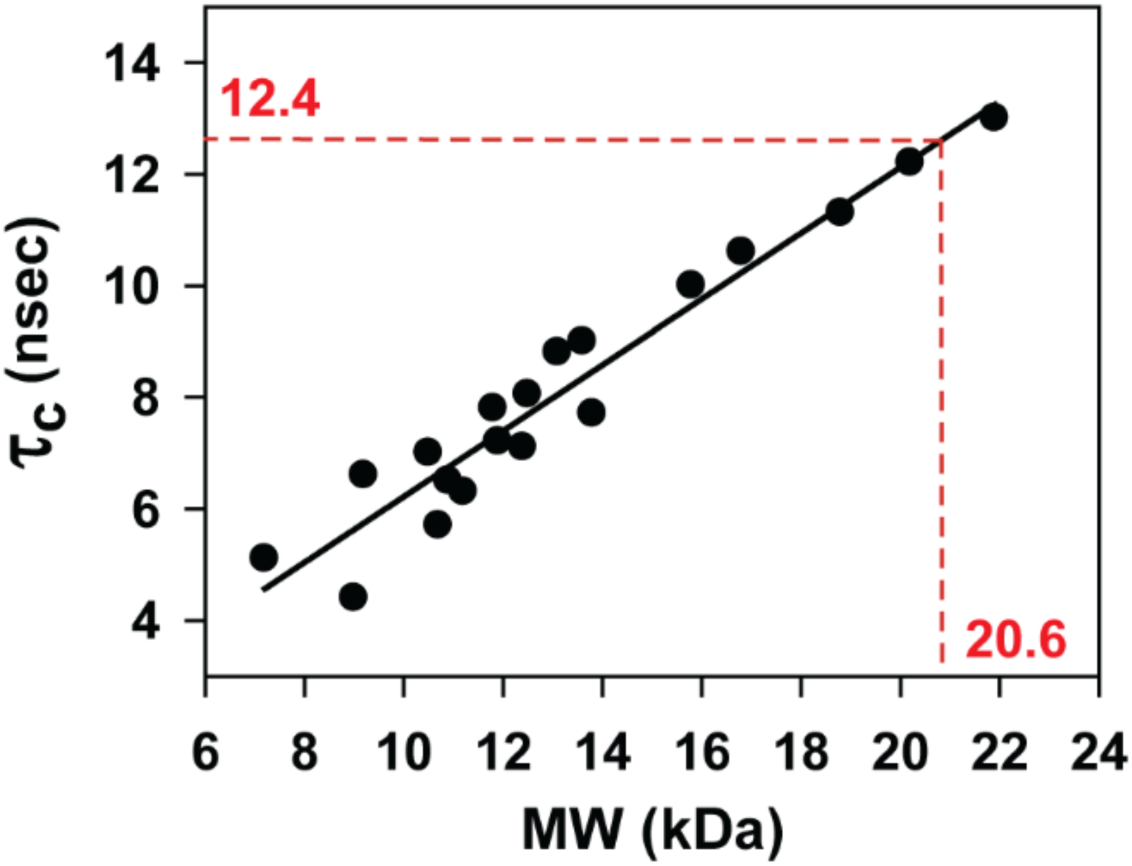
Standard curve of correlation time (τ_c_) versus molecular weight (MW). MW of SARS-CoV-2 Nsp7 at 298 K was estimated from its τ_c_ using linear regression analysis. The empirical data from NorthEast Structural Genomics consortium (NESG)^28^ were used to create the standard curve.

## Methods

### Sample Preparation and NMR Assignments

The ^13^C, ^15^N-labeled SARS-CoV-2 nsp7 sample was expressed and purified by the same methods as previously described in our NMR assignment paper.^20^ The recombinant *Escherichia coli* Rossetta2 pLysS was first grown in a M9 minimal medium containing 1 g/L ^15^N_4_HCl (Cambridge Isotope Laboratories), 4 g/L ^13^C_6_-D-Glucose (Cambridge Isotope Laboratories) and 100 mg/L ampicillin at 37 °C. 0.5 mM of IPTG was added when OD_600nm_ reached 0.8. The protein was expressed at 16 °C for 14-16 hours. After resuspending a cell pellet in buffer A (10 mM HEPES, pH 7.4, 30 mM imidazole and 2mM DTT), cells were lysed using a microfluidizer at 20,000 psi. The supernatant was filtered using a 0.45 μm vacuum filter before loading to a Ni-NTA agarose column (Qiagen). The protein was eluted with buffer B (buffer A with 300mM imidazole) and cleaved with 1% w/w TEV protease overnight at room temperature while dialyzing against 10 mM MOPS buffer (pH 7.0), containing 150 mM NaCl and 2 mM DTT. TEV protease and tag were removed via a second IMAC purification. The protein was further purified with the Superdex200 (GE Life Science) using the MOPS buffer used in the dialysis. The final NMR sample was prepared as 1.7 mM nsp7 containing 0.025 % NaN_3_ and 7 % D_2_O.

All NMR experiments were performed by 600 MHz Varian VNMRS and 600 MHz Bruker Avance III at 298 K. NMR data were acquired and processed using NMRPipe^30^ with SMILE^31^ for NUS reconstruction as described in the previous paper.^20^ Using pre-deposited chemical shift assignments (BMRB ID 50337),^20^ NOE data – NOESY ^1^H-^15^N HSQC, aliphatic NOESY ^1^H-^13^C HSQC, and aromatic NOESY ^1^H-13C HSQC – were assigned manually after automated NOE assignments with AUDANA algorithm^32^ in PONDEROSA-C/S software package^33^. NMRFAM-SPARKY^34^ in the virtual server of NMRbox35 was used for the assignments. The 3D NOESY spectra were aligned based on the chemical shift assignments using the *Align Spectrum* tool (the two-letter code *al*).

### Structure Determination

The structure of SARS-CoV-2 nsp7 at pH 7.0 was calculated using PONDEROSA-C/S software package.^33^ At first, the initial crude structure was obtained after iterative cycles of *AUDANA/AUDASA automation*. Distance and hydrogen bond constraints were automatically generated by AUDANA algorithm using the protein sequence, chemical shift assignments, and NOE data.^32^ Before submitting calculations, all unused resonances were removed by the two-letter code *dr*, and nomenclature and resonance deviations of chemical shift assignments were thoroughly verified using the *Check Protein Nomenclature* tool (*ca*) and the *Resonance Deviation Check* tool (*ck*) in NMRFAM-SPARKY, respectively. Chemical shifts assignments were also used to generate backbone dihedral angle constraints by TALOS-N.^36^ After every calculation, distance constraints were manually validated using the *Distance Constraint Validator* tool in Poky Analyzer,^37^ along with the manual NOE assignments. During the validations, all spectrum data and 3D atomic coordinates were visually examined. This was repeated until the number of long-range NOEs reached about 5 times of the number of residues.

The initial crude structure was refined by *Constraints-X only* option that uses simulated annealing and molecular dynamics protocols in Xplor-NIH.^38^ Similar to the previous step, distance and angle constraints were manually validated with each validator tool in Poky Analyzer. The structure went through several refinements until its 20 lowest models showed no mean NOE violations larger than 0.5 Å nor mean angle violations greater than 5°. The final structure of SARS-CoV-2 nsp7 was determined by *Final step with explicit H*_*2*_*O (x2)* option. Among 200 calculated structures, the 20 lowest energy models were selected after the molecular dynamics simulation of water solvation.^33^ The quality of the final structure was evaluated with Poky Analyzer and PSVS 1.5 software suite^39^ (Table 1). The Ramachandran plot analysis from PROCHECK^19^ revealed that 95.4 % of residues are in the most favored regions, followed by 3.0 % in additionally allowed regions, 1.6 % in generously allowed regions and 0.0 % in disallowed regions.

### Spin-relaxation Experiment

All spin-relaxation data were acquired on an Avance III Bruker spectrometer operating at 600MHz (^1^H) and equipped with a triple resonance cryogenic probe. The temperature of the sample was regulated at 298 K throughout the experiments. T_1_ (R_1_) experiment data were recorded as a series of interleaved 2D ^1^H-^15^N HSQC spectra using 12 relaxation delays, 80, 160, 240, 320, 400, 480, 560, 640, 800, 960, 1200 and 1520 ms. Similarly, T_2_ (R_2_) experiment data were acquired with 13 relaxation delays, 10, 20, 30, 40, 50, 60, 70, 80, 90, 100, 120, 140 and 160 ms. The recycle delay was set to 2.0sec for both T_1_ and T_2_ experiments. The hetNOE experiment data were obtained as two ^1^H-^15^N HSQC spectra from interleaving pulse sequences with (NOE) and without (no NOE) proton saturation. A 3s recycle delay was used for the no NOE experiment. This delay was replaced by a 3s proton saturation period in the NOE experiment. For all experiments, the offset and the spectral width were 4.76 and 16.7 ppm for the t_2_ dimension and 116.8 and 29.9 ppm for the t_1_ dimension, respectively.

Poky Notepad in the Poky software suite ^37^ was used to run scripts for calculating T_1_, R_1_, T_2_, R_2_, τ_c_, MW, and hetNOE. The decay curves of peak heights over relaxation delays were fitted using scipy.optimize.curve_fit ^40^, and the same fitting function was used to calculate T_1_ and T_2_ times, I(t) = I(0) Exp[(−1/T_1,2_) t], where I(t) is the peak intensity at a relaxation delay time (t). Similarly, R_1_ and R_2_ rates were calculated using the fitting function, I(t) = I(0) Exp[(-R_1,2_/1000) t]. The τ_c_ of SARS-CoV-2 nsp7 in the NMR sample was calculated as a function of the ratio of the average values of T_1_ and T_2_. ^28,29^ To minimize contributions from unfolded segments, only the peaks with ^1^H chemical shift higher than 8.5 ppm were considered for the τ_c_ calculation. The MW of the protein was approximated using the standard curve of τ_c_ versus MW created with empirical data from NESG^28^. Finally, the hetNOEs were calculated as the ratio of peak heights in the NOE spectrum with proton saturation and the reference spectrum without proton saturation. The errors of each hetNOE were estimated as [(σ_NOE_/I_NOE_)^2^ +(σ_Ref_/I_Ref_)^2^]^1/2^, where σ_NOE_ and σ_Ref_ represent the root-mean-square noises of background regions in each spectrum, and I_NOE_ and I_Ref_ are the intensities of a given peak. ^41^

## Acknowledgements

This research was funded by National Science Foundation DBI-2051595 and DBI-1902076, and the University of Colorado Denver [to W.L.]. This research was also supported by National Science Foundation EAGER MCB-2031269 [to K.H.W.] and the National Institutes for Health, National Institute for Allergy and Infectious Disease AI123498 [to R.N.K.]. This study utilized the National Magnetic Resonance Facility at Madison supported by NIH grant P41GM103399 (NIGMS), old number: P41RR002301. This study made use of NMRbox: National Center for Biomolecular NMR Data Processing and Analysis, a Biomedical Technology Research Resource (BTRR), which is supported by NIH grant P41GM111135 (NIGMS). Authors acknowledge the useful discussions in COVID19-NMR meetings.

## Author contributions statement

Y.L. structure, manuscript, analysis

M.T. NMR, manuscript

M.R. computational work

T.K.A., R.N.K. sample preparation

K.H.W., W.L. design, manuscript, direction

## Additional information

### Accession codes

Final coordinates of the solution structure of SARS-CoV-2 nsp7 at pH 7.0 are deposited in the PDB under the entry ID 7LHQ (BMRB ID: 50337).

## References

1 Harrison, A. G., Lin, T. & Wang, P. Mechanisms of SARS-CoV-2 Transmission and Pathogenesis. Trends Immunol 41, 1100–1115, doi:10.1016/j.it.2020.10.004 (2020).

2 Coronaviridae Study Group of the International Committee on Taxonomy of, V. The species Severe acute respiratory syndrome-related coronavirus: classifying 2019-nCoV and naming it SARS-CoV-2. Nat Microbiol 5, 536–544, doi:10.1038/s41564-020-0695-z (2020).

3 Zhou, P. et al. A pneumonia outbreak associated with a new coronavirus of probable bat origin. Nature 579, 270–273, doi:10.1038/s41586-020-2012-7 (2020).

4 Lu, R. et al. Genomic characterisation and epidemiology of 2019 novel coronavirus: implications for virus origins and receptor binding. Lancet 395, 565–574, doi:10.1016/S0140-6736(20)30251-8 (2020).

5 Perlman, S. & Netland, J. Coronaviruses post-SARS: update on replication and pathogenesis. Nat Rev Microbiol 7, 439–450, doi:10.1038/nrmicro2147 (2009).

6 Blanchard, E. & Roingeard, P. Virus-induced double-membrane vesicles. Cell Microbiol 17, 45–50, doi:10.1111/cmi.12372 (2015).

7 Wolff, G., Melia, C. E., Snijder, E. J. & Barcena, M. Double-Membrane Vesicles as Platforms for Viral Replication. Trends Microbiol 28, 1022–1033, doi:10.1016/j.tim.2020.05.009 (2020).

8 Wu, H. Y. & Brian, D. A. Subgenomic messenger RNA amplification in coronaviruses. Proc Natl Acad Sci U S A 107, 12257–12262, doi:10.1073/pnas.1000378107 (2010).

9 Sicari, D., Chatziioannou, A., Koutsandreas, T., Sitia, R. & Chevet, E. Role of the early secretory pathway in SARS-CoV-2 infection. J Cell Biol 219, doi:10.1083/jcb.202006005 (2020).

10 Song, J. et al. Mapping the Nonstructural Protein Interaction Network of Porcine Reproductive and Respiratory Syndrome Virus. J Virol 92, doi:10.1128/JVI.01112-18 (2018).

11 Kirchdoerfer, R. N. & Ward, A. B. Structure of the SARS-CoV nsp12 polymerase bound to nsp7 and nsp8 cofactors. Nat Commun 10, 2342, doi:10.1038/s41467-019-10280-3 (2019).

12 Gao, Y. et al. Structure of the RNA-dependent RNA polymerase from COVID-19 virus. Science 368, 779–782, doi:10.1126/science.abb7498 (2020).

13 Hillen, H. S. et al. Structure of replicating SARS-CoV-2 polymerase. Nature 584, 154–156, doi:10.1038/s41586-020-2368-8 (2020).

14 Subissi, L. et al. One severe acute respiratory syndrome coronavirus protein complex integrates processive RNA polymerase and exonuclease activities. Proc Natl Acad Sci U S A 111, E3900–3909, doi:10.1073/pnas.1323705111 (2014).

15 Peng, Q. et al. Structural and Biochemical Characterization of the nsp12-nsp7-nsp8 Core Polymerase Complex from SARS-CoV-2. Cell Rep 31, 107774, doi:10.1016/j.celrep.2020.107774 (2020).

16 Wang, Q. et al. Structural Basis for RNA Replication by the SARS-CoV-2 Polymerase. Cell 182, 417–428 e413, doi:10.1016/j.cell.2020.05.034 (2020).

17 Peti, W. et al. Structural genomics of the severe acute respiratory syndrome coronavirus: nuclear magnetic resonance structure of the protein nsP7. J Virol 79, 12905–12913, doi:10.1128/JVI.79.20.12905-12913.2005 (2005).

18 Johnson, M. A., Jaudzems, K. & Wuthrich, K. NMR Structure of the SARS-CoV Nonstructural Protein 7 in Solution at pH 6.5. J Mol Biol 402, 619–628, doi:10.1016/j.jmb.2010.07.043 (2010).

19 Laskowski, R. A., Rullmannn, J. A., MacArthur, M. W., Kaptein, R. & Thornton, J. M. AQUA and PROCHECK-NMR: programs for checking the quality of protein structures solved by NMR. J Biomol NMR 8, 477–486, doi:10.1007/BF00228148 (1996).

20 Tonelli, M., Rienstra, C., Anderson, T. K., Kirchdoerfer, R. & Henzler-Wildman, K. 1H, 13C, and 15N backbone and side chain chemical shift assignments of the SARS-CoV-2 non-structural protein 7. Biomol NMR Assign, doi:10.1007/s12104-020-09985-0 (2020).

21 Buckley, J. T. et al. Protonation of histidine-132 promotes oligomerization of the channel-forming toxin aerolysin. Biochemistry 34, 16450–16455, doi:10.1021/bi00050a028 (1995).

22 Dai, Z., Kim, J. H., Tonelli, M., Ali, I. K. & Markley, J. L. pH-induced conformational change of IscU at low pH correlates with protonation/deprotonation of two conserved histidine residues. Biochemistry 53, 5290–5297, doi:10.1021/bi500313t (2014).

23 Homeyer, N., Essigke, T., Ullmann, G. M. & Sticht, H. Effects of histidine protonation and phosphorylation on histidine-containing phosphocarrier protein structure, dynamics, and physicochemical properties. Biochemistry 46, 12314–12326, doi:10.1021/bi701228g (2007).

24 Langella, E., Improta, R. & Barone, V. Checking the pH-induced conformational transition of prion protein by molecular dynamics simulations: effect of protonation of histidine residues. Biophys J 87, 3623–3632, doi:10.1529/biophysj.104.043448 (2004).

25 Medina, E., Villalobos, P., Conuecar, R., Ramirez-Sarmiento, C. A. & Babul, J. The protonation state of an evolutionarily conserved histidine modulates domainswapping stability of FoxP1. Sci Rep 9, 5441, doi:10.1038/s41598-019-41819-5 (2019).

26 Mueller, D. S. et al. Histidine protonation and the activation of viral fusion proteins. Biochem Soc Trans 36, 43–45, doi:10.1042/BST0360043 (2008).

27 Paroutis, P., Touret, N. & Grinstein, S. The pH of the secretory pathway: measurement, determinants, and regulation. Physiology (Bethesda) 19, 207–215, doi:10.1152/physiol.00005.2004 (2004).

28 Rossi, P. et al. A microscale protein NMR sample screening pipeline. J Biomol NMR 46, 11–22, doi:10.1007/s10858-009-9386-z (2010).

29 Kay, L. E., Torchia, D. A. & Bax, A. Backbone dynamics of proteins as studied by 15N inverse detected heteronuclear NMR spectroscopy: application to staphylococcal nuclease. Biochemistry 28, 8972–8979, doi:10.1021/bi00449a003 (1989).

30 Delaglio, F. et al. NMRPipe: a multidimensional spectral processing system based on UNIX pipes. J Biomol NMR 6, 277–293, doi:10.1007/BF00197809 (1995).

31 Ying, J., Delaglio, F., Torchia, D. A. & Bax, A. Sparse multidimensional iterative lineshape-enhanced (SMILE) reconstruction of both non-uniformly sampled and conventional NMR data. J Biomol NMR 68, 101–118, doi:10.1007/s10858-016-0072-7 (2017).

32 Lee, W., Petit, C. M., Cornilescu, G., Stark, J. L. & Markley, J. L. The AUDANA algorithm for automated protein 3D structure determination from NMR NOE data. J Biomol NMR 65, 51–57, doi:10.1007/s10858-016-0036-y (2016).

33 Lee, W., Stark, J. L. & Markley, J. L. PONDEROSA-C/S: client-server based software package for automated protein 3D structure determination. J Biomol NMR 60, 73–75, doi:10.1007/s10858-014-9855-x (2014).

34 Lee, W., Tonelli, M. & Markley, J. L. NMRFAM-SPARKY: enhanced software for biomolecular NMR spectroscopy. Bioinformatics 31, 1325–1327, doi:10.1093/bioinformatics/btu830 (2015).

35 Maciejewski, M. W. et al. NMRbox: A Resource for Biomolecular NMR Computation. Biophys J 112, 1529–1534, doi:10.1016/j.bpj.2017.03.011 (2017).

36 Shen, Y. & Bax, A. Protein structural information derived from NMR chemical shift with the neural network program TALOS-N. Methods Mol Biol 1260, 17–32, doi:10.1007/978-1-4939-2239-0_2 (2015).

37 Lee, W., Rahimi, M., Lee, Y. & Chiu, A. POKY: a software suite for multidimensional NMR and 3D structure calculation of biomolecules. Bioinformatics, doi:10.1093/bioinformatics/btab180 (2021).

38 Schwieters, C. D., Bermejo, G. A. & Clore, G. M. Xplor-NIH for molecular structure determination from NMR and other data sources. Protein Sci 27, 26–40, doi:10.1002/pro.3248 (2018).

39 Bhattacharya, A., Tejero, R. & Montelione, G. T. Evaluating protein structures determined by structural genomics consortia. Proteins 66, 778–795, doi:10.1002/prot.21165 (2007).

40 Virtanen, P. et al. SciPy 1.0: fundamental algorithms for scientific computing in Python. Nat Methods 17, 261–272, doi:10.1038/s41592-019-0686-2 (2020).

41 Farrow, N. A. et al. Backbone dynamics of a free and phosphopeptide-complexed Src homology 2 domain studied by 15N NMR relaxation. Biochemistry 33, 5984–6003, doi:10.1021/bi00185a040 (1994).

